# A Computational Framework for DNA Sequencing-Based Microscopy

**DOI:** 10.1101/476200

**Authors:** Ian T. Hoffecker, Yunshi Yang, Giulio Bernardinelli, Pekka Orponen, Björn Högberg

## Abstract

Barcoded DNA polony amplification techniques provide a means to impart a unique sequence identity onto specific locations of a surface wafer or chip. We describe a method whereby micro-scale spatial information such as the relative positions of biomolecules on a surface can be transferred to a sequence-based format and reconstructed into images without the use of conventional optics. The principle is based on the pair-wise association of uniquely tagged and spatially adjacenct polonies. The network of polonies connected by shared borders forms a graph whose topology can be reconstructed from a set of edges derived from pairs of barcodes associated with each other during a polony crosslinking phase, the sequences of which could be determined by isolation of the DNA and next-gen sequencing. We developed a mathematical and computational framework for this principle called Polony Adjacency Reconstruction for Spatial Inference and Topology and show that Euclidean spatial data may be partially stored and transmitted in the form of untethered graph topology. This effect may be exploited to form images by transferring molecular information from a surface of interest, which we demonstrated *in silico* by reconstructing images formed from stochastic transfer of hypothetical red, green, and blue molecular markers. The theory developed here could serve as a guide for a highly automated, multiplexable, and potentially super resolution imaging method based on molecular information encoding and transmission.

## 1 Introduction

### 1.1 Transfer of Spatial Information to DNA Sequences

Small-scale imaging techniques have, from their beginnings to modern techniques like super resolution microscopy [7, 3, 16], relied on optics to amplify and enlarge signals derived from initially confined spatial regions. Notable exceptions include atomic force microscopy and transmission electron microscopy which achieve resolutions superior to optical methods by utilizing effectively smaller probes to interact with the sample e.g. the wavelength of an electron is smaller than that of light as are the minute interactions between an AFM tip and sample surface. DNA has a high information density, with storage levels of 5.5 petabits per cubic millimeter achieved [4], making it an attractive medium for encoding spatial information at micro-to nano-scales. In this paper, we present a theoretical foundation for a spatial information encoding approach that utilizes DNA sequencing and graph theory that in principle could be used to generate statistical maps of biomolecules at micro- and nano-scales as well as whole images.

**Fig. 1.**
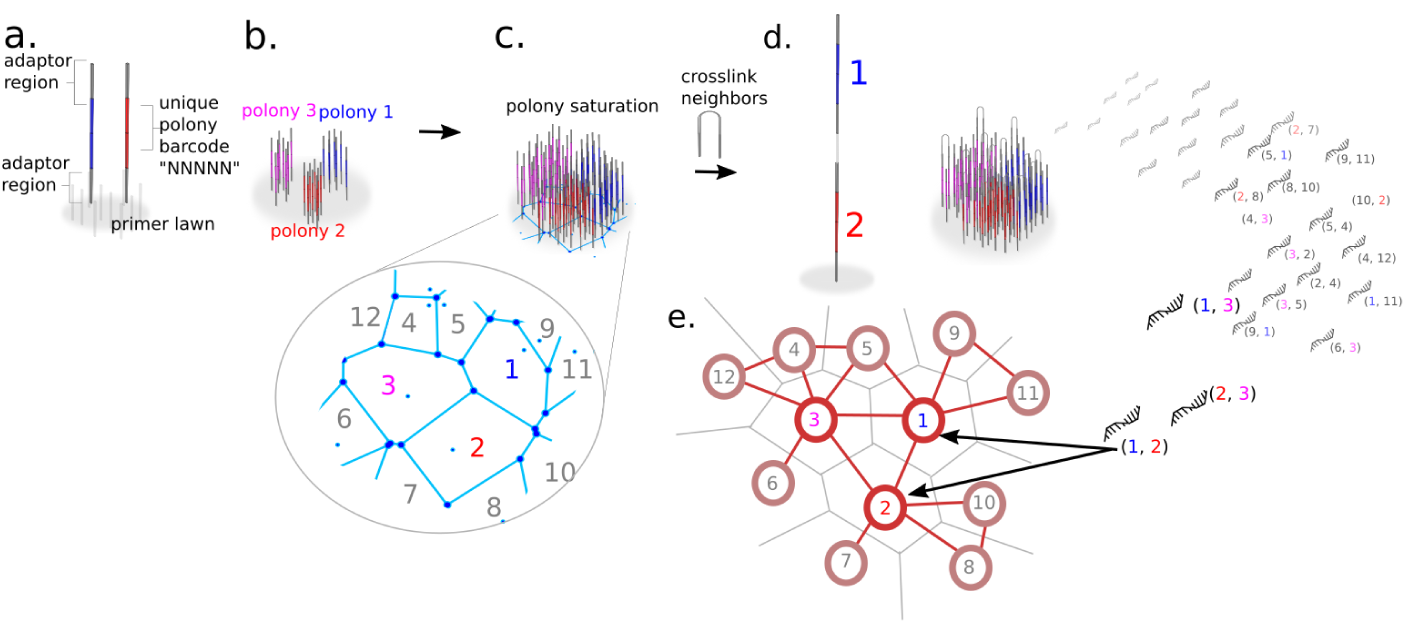
Encoding and recovering metrics through polony adjacency. (A) Seed molecules with a unique barcode sequence land randomly on a surface of primers. (B) Local amplification of seed molecules produces sequence-distinct polonies. (C) Saturation of polonies occurs when polonies are blocked from further growth by encountering adjacent polonies, forming a tessellated surface. (D) Random crosslinking of adjacent strands leads to pairwise association of nearby barcodes. (E) Recovery and sequencing of barcode pairs is used to reconstruct a network with similar relative positions of polonies as the original surface.

DNA has been used in conjunction with optical techniques to convey spatial information such as with proximity ligation assay (PLA) [18], and DNA PAINT [8] where DNA plays a role as a signifier of molecular interactions by conveying this information through fluorescent signals and optical readout/detection of binding events. This principle also underlies a family of techniques for connecting spatial locations with single cell RNA sequencing data. In these techniques, recovery of spatial locations are important for distinguishing tissue types and subtle patterns in expression of cells varying across distances, information that would otherwise be lost during the dissociation of cells prior to single cell sequencing. The operating principle is to use *a priori* knowledge of spatial marker genes to associate unknown genes to approximate locations, the *a priori* data is in most cases obtained with microscopy such as with *in situ* hybridization or mathematical modelling of spatial expression patterns to retrieve locations of associated genes following single cell sequencing[21, 9, 17, 1, 6, 12]. Alternatively, direct microscopy-based *in situ* sequencing methods have emerged in order to achieve precise context-sensitive spatial transcriptomic information without needing to scramble spatial data by dissociation prior to sequencing[22].

Next generation sequencing strategies have made the rapid and large scale reading of information stored in DNA an efficient process. However there is an absence of strategies for encoding spatial information in a way that is preserved during the scrambling during isolation and recovery from *in situ* contexts that can then be read and recovered with sequencing. A few techniques achieve this goal by encoding spatial information directly into the molecular format, e.g. in the form of DNA read during sequencing along with transcriptomic data. These methods are based on artificial generation of an addressable surface using printing or lithographic means similar to that employed for microarray manufacture [5, 19].

Herein, we describe a computational framework for a method called Polony Adjacency Reconstruction for Spatial Inference and Topology (PARSIFT), which is designed to accomplish the encoding of spatial information, for example the positions of specific molecules relative to others in a 2D plane, directly into a DNA-based format without transduction of information through any other medium. The principle is to achieve this without *a priori* surface addressing, circumventing the need for printing strategies. Instead PARSIFT utilizes topological information, i.e. the connectivity of vertices in a graph of spatially related DNA sequences, as a means to partially preserve Euclidean spatial information, and next-gen sequencing as a means to recover that information by post processing.

Encoding of topological data in DNA sequence format is possible by using DNA barcodes also known as unique molecular identifiers (UMI’s), i.e. randomly generated sequences of synthetic DNA. Barcodes that are associated with spatial patches can establish an identity for those locations, each patch distinguishable from another based on its sequence. A DNA barcode with 10 bases has 1.04e6 possible sequences, and larger barcodes can be used to create effectively unique labels in a system. The basic unit of topological data is an edge or association between two adjacent patches by physically linking between their respective barcodes. Topological mapping with barcoding has been used to map neural connectomes by the strategy of assembling a network from cells that share common barcodes transmitted by viruses able to traverse cells and leave a unique sequence identifier with each visit [10, 14, 15].

The assignment of barcodes to patches of a surface can be accomplished with surface polony generation methods such as bridge amplification [2], a 2-primer rolling circle amplification-based method [11], template walking amplification [13], whereby unique “seed” strands are captured by the surface containing primer strands (Figure 1 A) and then locally amplified in the immediate spatial vicinity of where they landed thus generating a diverse surface of distinct patches of amplified DNA (Figure 1 B). A polony is defined as one of these locally amplified patches of DNA derived from a single seed molecule.

By growing polonies on a surface of primers to the point of saturation (Figure 1 C), i.e. a point where nearby primers are depleted and growing polonies grow to encounter the boundaries of other adjacent polonies, a tessellation of neighboring polonies is created. Each member of the tessellation, a polony, has a limited number of immediately adjacent neighboring polonies with distinct sequence due to the use of barcodes in each of the seed strands. Though each patch is associated with a unique sequence according to its parent seed molecule, isolation of this DNA and subsequent sequencing would scramble information about the polony’s position or its neighboring polonies. Thus the critical step is to crosslink strands from each polony to strands from adjacent polonies (Figure 1 D) in a way that enables both barcodes to be sequenced together in a single read. Recovery of the strands, i.e. stripping them from the surface followed by next-gen sequencing would then preserve topological association of neighboring polonies in the form of covalently linked pairs of barcodes - a complete set of which might be used to completely reconstruct the topological network of adjacent polonies without direct knowledge of their original coordinates (Figure 1 E). For seed distributions without long-range systematic variation, i.e. those that are Poisson distributed, we show that topological information alone, constrained by being a 2D planar network with known boundary geometry, retains significant spatial metrics with only local and boundary-related distortions that become insignificant at scales greater than those on the order of individual polonies. By generating such a mappable surface, we suggest that spatial analysis of other molecules could be accomplished by covalent association with the polony surface, enabling inference of molecular spatial distributions and construction of images where polonies serve as pixels. At the time of this manuscript’s publication online as a preprint, we are aware of a preprint by Weinstein, Regev, and Zhang that was made available immediately prior whose contribution is complementary to ours, describing a method of DNA-based microscopic image formation on the basis of UMI diffusion and concatenation and reconstruction of a large image by assembling local connectivity [23].

## 2 Theoretical Framework

### 2.1 Voronoi Tessellation as a Model of Polony Saturation

The spatial distribution of polonies on a surface, the *a priori* Euclidean information that is not explicitly accessible after isolation, can in principle be preserved by associations between adjacent polony sequences and recovered with sequencing. Information that is available after sequencing and subsequent transformations of that data are then referred to as *a posteriori*.

Assume that seed molecule amplification takes the form of uniform circular growth. At the point of saturation polonies have amplified to the extent that expanding boundaries are restricted from further growth by having encountered neighboring expanding polonies, and the set of boundaries forms a Voronoi tessellation *S*. Let the surface be represented with *X*, a real coordinate plane in 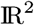 with a Euclidean distance function *d*(*x*_*i*_, *x*_*j*_)∀*x* ∈ *X*. Let *K* be a set of indices and let *p*_*k*(*k*∈*K*)_ ∈ *P* be a tuple in the space *X* residing within a circle centered at an origin *x*_0_ with radius *a* and Poisson distributed with spatial frequency *λ*.

At the point of saturation, each polony is represented by a facet *s*_*k*_ ∈ *S* or the set of points closest to its corresponding seed molecule *p*_*k*_.

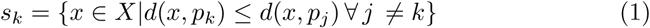

Every Voronoi tessellation has a corresponding Delaunay triangulation *D* = (*P, E*), a triangulation of the set of points *P* and set of edges *E* satisfying the empty circle property: no circumcircle of any triangle in *D* has a point in its interior. We refer to the graph, defined by its set of vertices and edges, without Euclidean metrics as the *untethered graph*. Figure 2 A shows an example Voronoi tessellation formed from a set of 9 seed points that fall within a circular region. Figure 2 B shows the Delaunay triangulation corresponding to 2 A, and the Delaunay-derived, non-embedded untethered graph *G* = (*K, E*) which is the set of Delaunay edges and vertices lacking coordinates. The untethered graph retains its topological metrics, i.e. a geodesic distance function defined as the fewest number of edges that must be traversed between two vertices. A geodesic distance 1 vertex, defined in relation to another vertex of distance 0 (origin), is one whose shortest path is 1 edge. A distance 2 vertex is one whose geodesic distance is 2 edges from the distance 0 vertex, and so on.

**Fig. 2.**
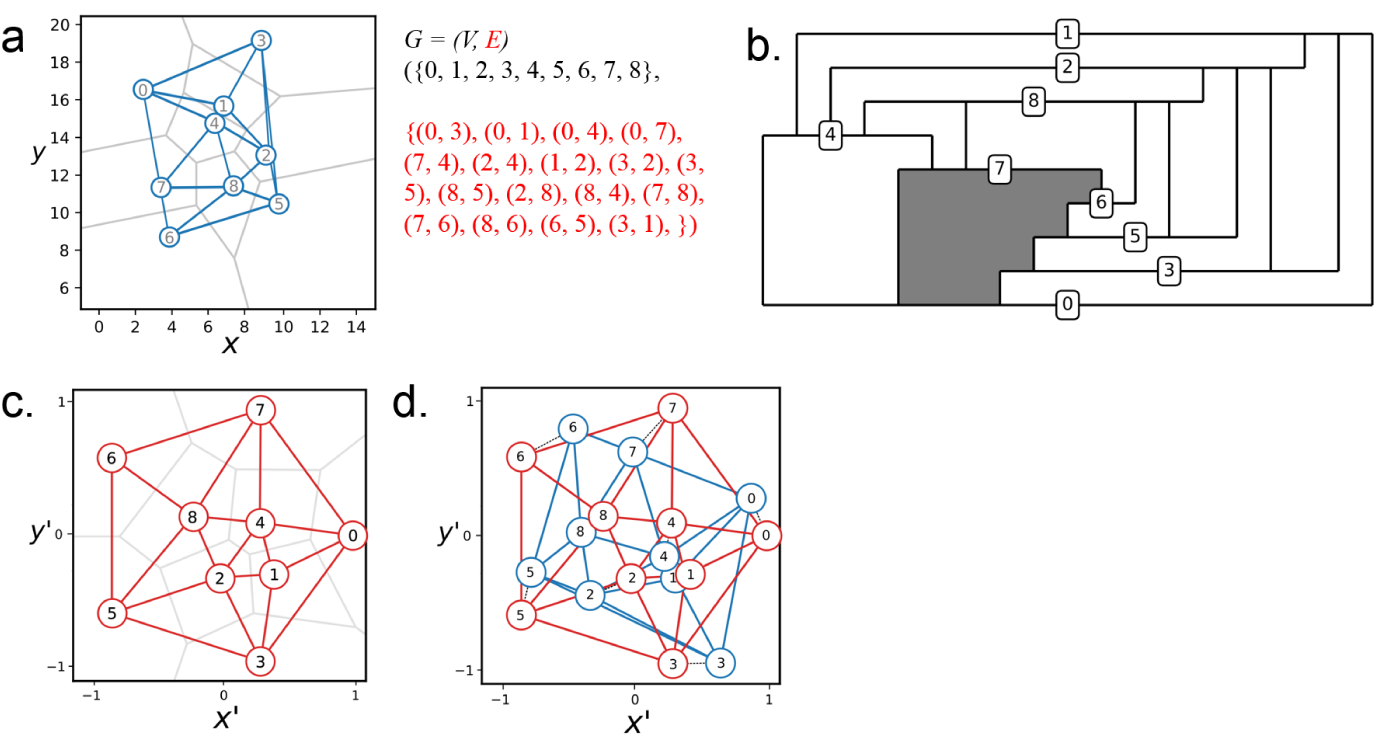
Encoding and recovering metrics via topology. (A) Seed molecule points distributed randomly on a real coordinate plane and used to compute a Voronoi tessellation (blue lines) denoting the boundaries of polonies. Corresponding Voronoi tessellation is shown in gray. (B) Planar embedding diagram with numbered horizontal lines denoting vertices and vertical lines denoting edges. Filled in area shows the face with the most edges. (C) Tutte embedding constructed from the graph from A by arranging the largest face uniformly on the unit circle. (D) Alignment of Tutte embedding from C with the original Delaunay from A with dotted black lines drawn between corresponding points.

An individual polony is identified by its barcode, a sequence of the alphabet *Σ* = *A, T, G, C* . We represent a barcode *w* as a string of length |*w*| = *l* with each letter ∈ *Σ*. The set of all *n* possible barcode sequences of length *l* is *Σ*^l^. Let *W*_*A*_ ⊆ *Σ*^*l*^ be a set of barcodes representing all polonies enclosed within a circular region of *S* called *A*. The average number of polonies *m* enclosed by *A* is thus *Aλ*.

The probability of *m* polonies all having unique barcodes for a sequence of length *l* can be represented with binomial coefficients as follows:

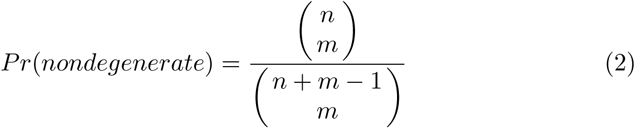

or approximated with the series:

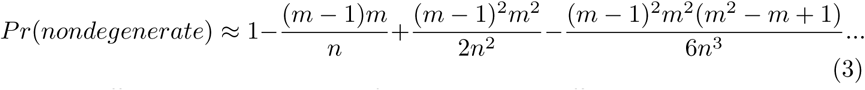

For sufficiently large number of barcodes *n* or sufficiently small *m* the number of possible sequences is much greater than the number of polonies. For a 20 nucleotide barcode and a region with 4000 polonies and assuming a naive, uniformly distributed, base composition, the probability of them all being unique is *p* ≈ 0.99998. We thus consider the ideal scenario where *n* ≫ *m*, and each polony *s*_*k*_ is uniquely distinguishable by its barcode sequence *w*_*k*_. Thus, the existence of a barcode pair {*w*_*i*_, *w*_*j*_} implies physical adjacency of two polonies between which crosslinks have associated their unique identifiers.

### 2.2 Geodesic Metrics as a Proxy for Euclidean Metrics

The untethered graph *G* = (*V, E*) may be constructed from the set of all edges

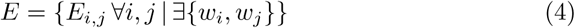

Through the Delaunay triangulation, a set of topological relationships are defined based on spatial determinants, creating a connection between Euclidean and geodesic metrics. Thus if we can robustly identify topological relationships by sequencing pairs of barcodes, we could recover Euclidean metrics by embedding the points in a plane and determining spatial positions of each vertex that obey as much as possible the known constraints of the original metrics. Given a known boundary geometry, e.g. a circle with known radius *a*, and knoweldge that polonies are Poisson-distributed, we conjecture that for any two vertices in the non-embedded graph *G* with a *N* length geodesic shortest path distance, there exists a vertex along that shortest path with *N* − 1 geodesic distance that is also closer to the origin in *Euclidean* distance.

*Conjecture 1.* Let {*k*_0_, *k*_*N*_, *k*_*N* − 1_} ⊆ *K* be any subset of three vertices in *K* that satisfies the property that *k*_*N*_ has at least one geodesic shortest path (i.e. hop count or number of edges) leading to a so-called origin *k*_0_ equal to *N* steps, and let *k*_*N* − 1_ be a vertex located on a geodesic shortest path to *k_N_* with its own geodesic shortest path to *k*_0_ equal to *N* − 1 steps. Let the set {*p*_*k*_, *p*_*k*_, *pk*_*N-1*_} ⊆ *P* be the corresponding set of real coordinate points in *D* in 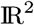. Then it is conjectured that

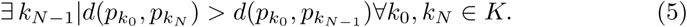

In other words, a graph derived from a Delaunay triangulation contains encoded in its geodesic topology a certain limited representation of the Euclidean spatial hierarchy that the Delaunay was generated from, whereby geodesic distance *n* vertices are further in Euclidean distance than at least one vertex lying on a shortest path at geodesic distance *n −* 1.

### 2.3 Embedding the Untethered Graph and Approximation of Spatial Position

By embedding the untethered graph in a Euclidean metric space, we can attempt to reconstruct the Euclidean hierarchy that is stored as a geodesic hierarchy. The embedding of *G* in a Euclidean metric space is defined by its graph and the positions of its vertices, so the embedded *a priori* graph is *{G, P }* = *D*. Our goal is to approximate *{G, P }* with an *a posteriori* embedding *{G, P′}*. We can define a new set of Euclidean tuples 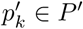 in a space *X*′ residing in a unit circle. The unit circle could be scaled according to a known boundary metric from the *a priori* seed distribution -i.e. if the area extracted was known to have a diamter of 10 *µm*. Each of the points 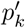 correspond to the vertices of *G*. Thus, together, the points *P′* and the edges *E* determine whether or not two edges cross each other in the embedded *G*. The goal is to determine the set of coordinates *P′* in a way that minimizes the distances between *a priori* and *a posteriori* points, i.e. 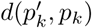). One approach is to find a set *P* that shares as many knowable *a posteriori* properties of *P* as possible.

One of these properties is planarity of the embedded graph {*G, P* }. This is due to the physical assumption that adjacency is determined by the nearest-neighborship of two polonies, and that an edge may not be formed if it must cross over a polony to bridge two non-neighboring polonies. Since the graph has no crosses and its points are embedded in a planar Euclidean space, the *D* and {*G, P*} are planar. A graph’s planar embedding can be determined algorithmically (Figure 2 C. The edges of this graph are taken from the set of edges produced from the Delaunay triangulation from Figure 2 A, scrambled and de-coupled from any explicit information about the *a priori* seed point distribution. Another criterion is that an average spatial density of the *a posteriori* vertex positions *λ′*should be obtained from the final distribution with no systematic variation across the reconstructed area. The points *P* should be consistent with that which could be drawn from a Poisson distribution.

A final property that should be satisfied is what we may call the same-Delaunay-topology criterion, in other words a new Delaunay triangulation *D′*generated from the *a posteriori* vertex positions *P′* should have the same geodesic structure *G′* as *G*. If *G* = *G′*, then *P′* shares in common with *P* all of the spatial constraints out of which *D* arose.

Since computing the positions that generate the same Delaunay topology as the original is computationally expensive, we developed an approach for approximating the positions with reasonable accuracy. A fast approximation of the original distribution can be obtained with a Tutte embedding or barycentric embedding [20], which takes a planar graph and forms a crossing-free straight-line embedding such that the outer face is a convex polygon and every interior vertex is located at the average (barycenter) of its neighboring vertex positions. If the outer face is fixed, the positions of the interior vertices are determined uniquely as the solution to a system of linear equations. The unique solution is always crossing-free, and every face is convex. The outer face can be found by enumerating all minimal faces and identifying that which has more than 3 edges, as all other faces in a true Delaunay graph are triangular.

We can make a conjecture analogous to Conjecture 1 about the coupling of Euclidean and geodesic metrics in the Tutte embedding, a suggestion that a Tutte embedding satisfies one of the major properties of the *a priori* distribution.

*Conjecture 2.* Let {*k*_0_, *k*_*N*_, *k*_*N*−1_} ⊆ *K* be any subset of three vertices in *K* that satisfies the property that *k*_*N*_ has at least one geodesic shortest path (i.e. hop count or number of edges) leading to a so-called origin *k*_0_ equal to *N* steps, and let *k*_*N − 1*_ be a vertex located on a geodesic shortest path to *k*_*N*_ with its own geodesic shortest path to *k*_0_ equal to *N* − 1 steps. Let the set 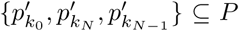 be a corresponding set of real coordinate points in the embedded graph *D′* in 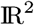. Then it is conjectured that

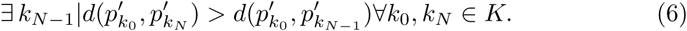

Or in other words, analogous to the spatial hierarchical relationship to geodesic hierarchy in the Delaunay triangulation, there is a corresponding spatial hierarchy in a Tutte embedding. We make this conjecture based on the observation that attempts to violate this rule by artificially placing a distance *N* vertex closer in Euclidean distance to some origin point than a distance *N*−1 vertex invariably result in the formation concave structures that are forbidden by the laws of the Tutte embedding. Another related observation is that the geodesic contour lines surrounding a vertex may never cross each other in Euclidean space. A geodesic contour *n* may not be closer at all points than some points of geodesic contour *n*−1, however there will always be some point on contour *n*−1 that is closer to the origin in Euclidean distance.

If spatial metrics about the original Euclidean boundary are known - for instance that we specify that points must lie within a circle, and the circle’s radius is known. Then the embedding may be scaled to match the original Euclidean metrics. Figure 2 C shows the Tutte embedding of untethered vertices from A-C with the outer face arranged uniformly around the unit circle. An alignment by scaling, rotating, translating, and flipping the a posteriori Tutte reconstruction of the seed positions to be aligned with the *a priori* Delaunay triangulation (Figure 2 C) shows that the basic spatial order is preserved, and vertices of the two graphs devate only locally.

## 3 Simulation and Generalization Beyond an Ideal (Triangulated) Posterior Graph

### 3.1 Simulation of Primer Lawn and Crosslinking

We performed an in silico proof of concept by simulating the random pairing of adjacent polony primer sites, scrambling of edge data, and reconstructing the untethered network based on topological information alone. We simulate the primer lawn with a hexagonal lattice, assuming the limit of molecular packing density Figure 3 A. Figure 3 B shows how crosslinking leads to random pairing of adjacent sites, some of which are self-pairing events (providing no additional information) and some of which are cross-polony sites that can be used to deduce the presence of a spatial boundary. The probabilistic nature of the pairing opens up the possibility to miss an existing boundary, particularly when the boundary is small. The untethered graph approaches the Delaunay triangulation at ideal pairing efficiency.

**Fig. 3.**
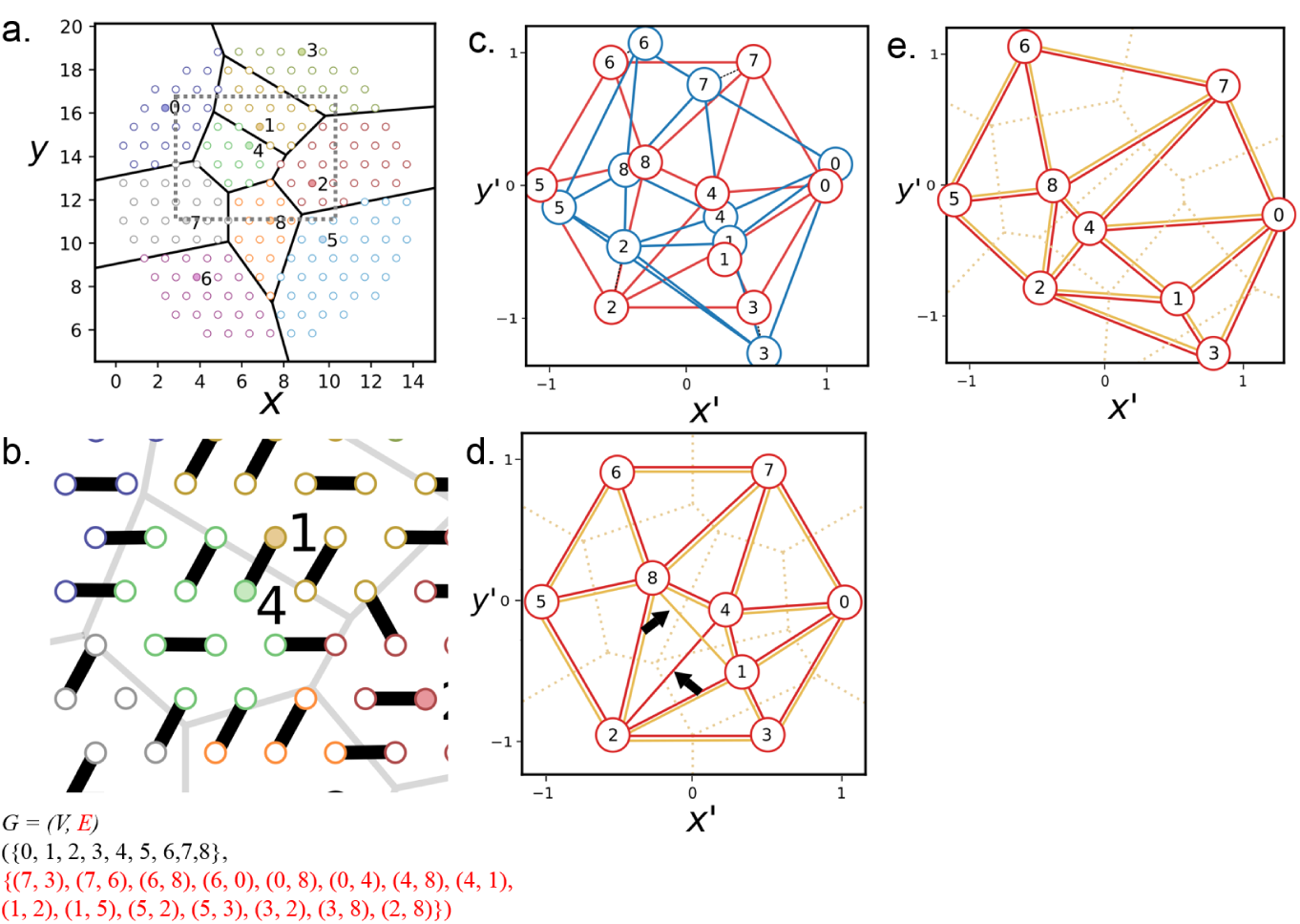
JIF and false discovery rate by subject area. Simulation of polony adjacency reconstruction. (A) Lattice diagram of primer lawn and polonies denoted with color and Voronoi face boundaries. (B) Illustration of random site pairing between adjacent primer sites, with those that bridge two polonies colored in red. The untethered graph (and the set of missed edges) is listed below, edges colored in red text. (C) Overlay of Tutte reconstruction from C with the Delaunay graph of the original Euclidean seed points. (D) Tutte reconstruction of the untethered graph formed from pairing events gathered from A. Re-computed Voronoi diagram is shown in dotted line. (E) Overlaid re-computed Delaunay-derived graph and embedded graph after adjustment for matching Delaunay topology

### 3.2 Reconstruction by Tutte Embedding

The edges formed from simulated cross-linking were scrambled, i.e. decoupled from Euclidean coordinates of the *a priori* seed distribution. We reconstructed the topological network from the list of edges and performed planar embedding, outer face identification, and Tutte embedding as with the ideal Delaunay case described in Figure 2. In practice, one does not have access to the original Euclidean seed positions, however for the purpose of characterizing distortion, we perform an alignment of the reconstructed graph with the original Delaunay graph (3 C) for the Tutte embedding approach and spring relaxation approach respectively. This is done by isotropically scaling, rotating, translating, and flipping the planar graph to minimize the distance between corresponding vertices by stochastic gradient descent. We can see that relative positions are preserved albeit with local distortion that leads to slight displacement of each reconstructed vertex relative to its original seed counterpart.

For relatively small graphs we could perform iterative adjustment to the positions, initialized with the Tutte embedding, until they form a Delaunay with the same topology as the original Delaunay. This was accomplished with a simulated annealing whereby with each iteration, the Delaunay graph generated by the current set of Euclidean coordinates (Figure 3 D orange) is compared with the untethered graph (Figure 3 D red), its topology derived from the Delaunay triangulation of the *a priori* seed distribution. After adjustment, a final graph satisfies the property that its Delaunay triangulation has the same topology as the untethered graph (Figure 3 E). The a posteriori positions that satisfy this same-Delaunay topology criterion thus satisfy all of the constraints of the original Delaunay triangulation.

### 3.3 Stamping and Image Formation

Approximate spatial location of each polony may be transmitted through this means. This could then be exploited to provide spatial information about objects of interest by association with polonies that we can trace the location of by PARSIFT. We devised a basic model of image reconstruction from the principle of contact-based transfer of molecules of interest to the mapped surface, i.e. a kind of molecular stamp. As proof of concept, we use an image (Figure 4 A) as a representation of a hypothetical probability distribution of 3 types of molecular markers (red, green, and blue). The image represents a surface of interest that we would like to sample from, for example a cell surface covered in oligo-tagged antibodies that would associate covalently with the mapped polony surface (Figure 4 B). The darkness and color of the image corresponds to the density of such markers and thus the probability that a marker of a particular color is placed on the polony surface. To simulate contact based transfer of the surface of interest with the polony surface, the overlaid lattice of primer sites denotes points where a Monte Carlo sampling will occur in the corresponding position in the image. If the image pixel at a given primer site location has an RGB value dominated by red and green for example, then there is a higher probability of that site being occupied by either a green or red marker (Figure 4 C). Since no spatial information is retained below the resolution of a single Voronoi face, the positions of markers within a polony is scrambled. The final rgb value of the face can be determined by tallying the markers that have associated with the primer sites in the polony as well as the number of un-associated sites (Figure 4 D). In this way we form a pixel from each polony/Voronoi face from which a complete image may be constructed by the reconstruction procedure described above.

**Fig. 4.**
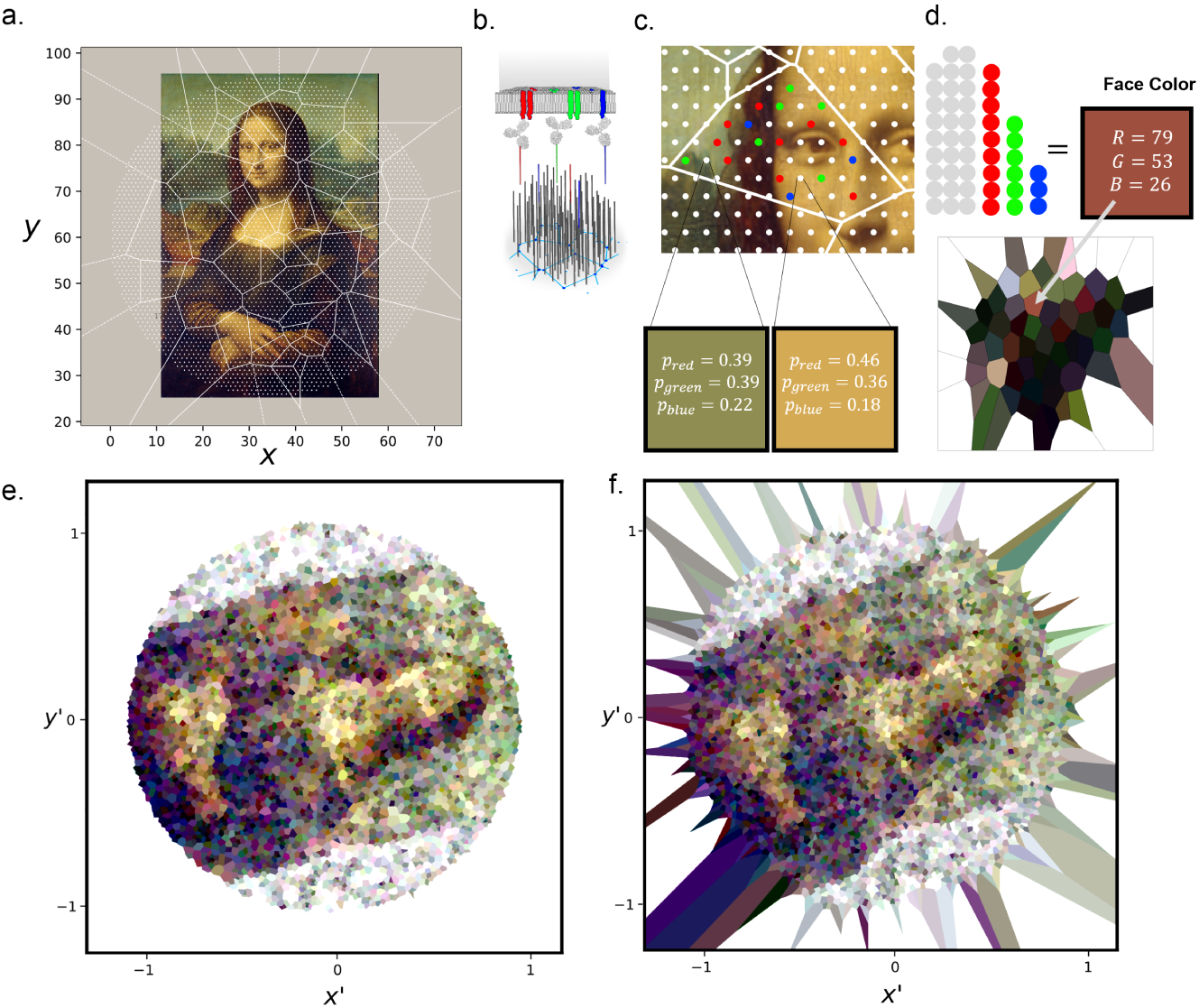
Transfer of surface of interest to the mappable polony surface for image reconstruction. (A) An image is overlaid on a simulated mesh with 65 polonies. (B) Illustrated concept of molecular markers representing 3 different colors/targets transferred by contact to the polony surface where they are covalently incorporated and associated with one of the polony barcodes. (C) Monte Carlo sampling procedure to determine whether a marker is associated with a given primer site and if so which color by taking the probability from the RGB value normalized to 1 at the corresponding position in the image. (D) Tallying of markers and empty sites within a given polony/Voronoi face is then used to determine the color of that ”pixel” and a 65-pixel image (lower pane) is formed by coloring each face accordingly. (E) Larger scale reconstruction using the Tutte embedding approach of image from A but with 4000 polonies. (F) Reconstruction analogous to F but with the spring relaxation approach.

Figure 4 E shows a Tutte reconstruction with 4000 polonies and a lattice density with an average 125.7 primer sites per polony. Note how rotational information is not preserved from the original image, however structure and features are represented from the original. Figure 4 F shows a reconstruction with the same parameters as E but reconstructed using the non-deterministic spring relaxation based procedure to generate the final positions. Note how the average size of polony is more uniform in the case of F compared to E which has several enlarged and contracted polonies, a distortion due to the combination of randomly missed edges and Tutte reconstruction which tends to expand gaps in the mesh.

## 4 Assessment of Distortion and Precision

Distortions are limited to local scales. We looked at the alignment errors of each point in the a posteriori reconstructed distribution compared to the original *a priori* seed distribution to obtain an average error normalized to the unit circle, i.e. with a value of 1.0 corresponding to the circle radius. This was done for multiple numbers of polonies and 4 different lattice densities for the Tutte embedding approach (Figure 5 A) as well as spring relaxation (Figure 5 B). In both cases we see that relative error is higher when there are fewer polonies but that this value decays gradually with more polonies. We can visualize the distortion (Figures 5 C and D) with lines drawn between corresponding vertices in the a posteriori reconstruction and the *a priori* seed distribution, and plotted as a scatter as a function of radius extending from the center of the reconstructed area. We see that while there are some systematic distortions at the local scale mostly near the boundaries, such distortions do not extend further than a few polonies. From this we can conclude that this approach should be effective for determining global structural properties so long as constraints of the simulation are met such as Poisson distributed seed points, uniform sampling from the stamped surface of interest, and uniform boundary geometry.

**Fig. 5.**
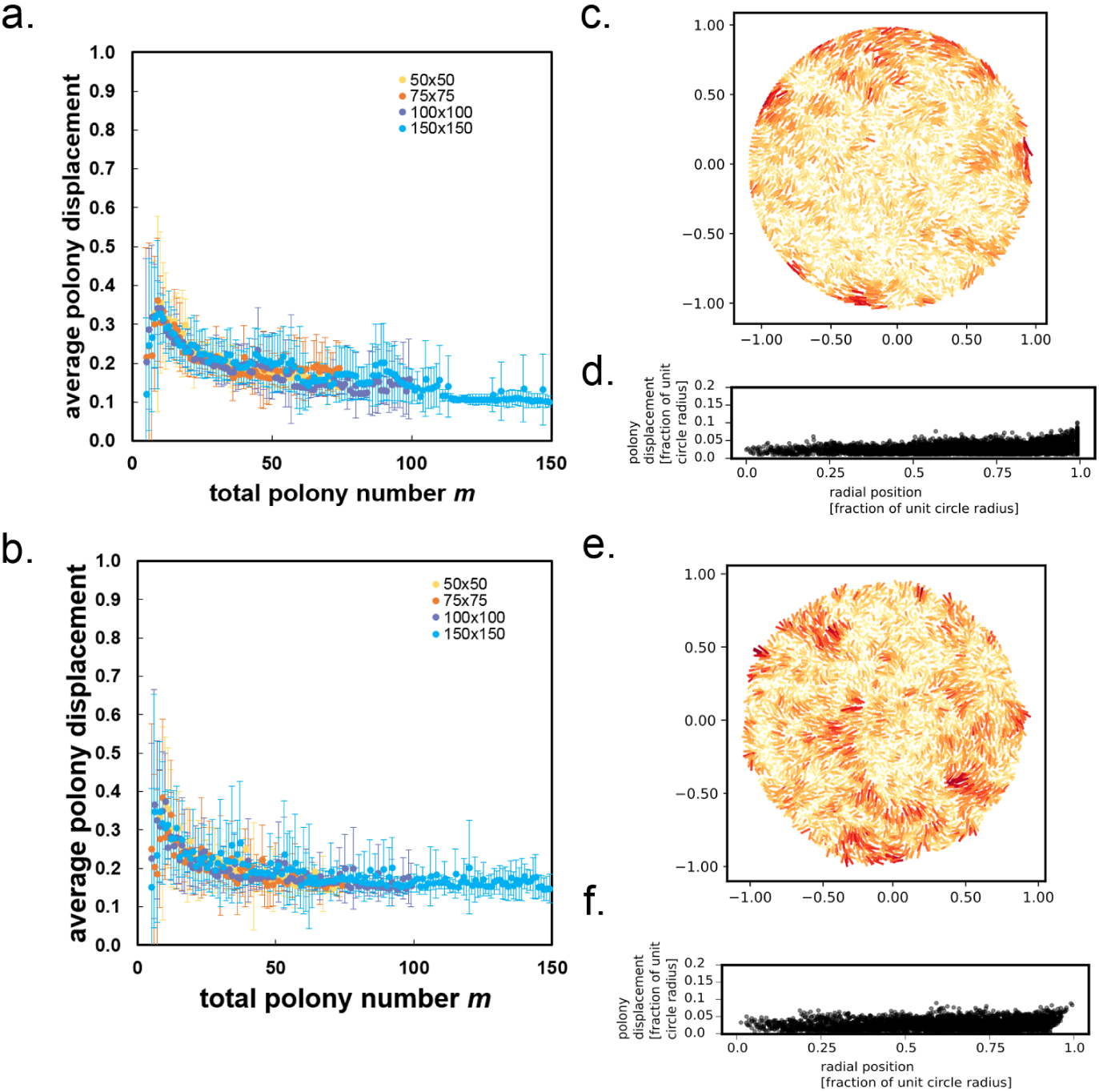
Distortion estimation from *a priori* and *a posteriori* vertex position comparison. (A) The average distance between a vertex *a priori* and *a posteriori* position with region normalized to unit circle, and varied for different numbers of polonies and primer site lattice densities, for the Tutte embedding reconstruction approach. (B) The average distance between a vertex *a priori* and *a posteriori* position with region normalized to unit circle, and varied for different numbers of polonies and primer site lattice densities, for the spring relaxation reconstruction approach. (C) Visualization of distortion of a 4000 polony *a posteriori* Tutte embedding aligned with the *a priori* seed positions. Lines a drawn between corresponding vertices with color map indicative of line length (red being longer). (D) Radial profile of Tutte embedding reconstruction from C Scatter plot of *a posteriori-a priori* distances as a function of radius from the center of the reconstructed area. Red line shows a moving average (10 point). (E) Distortion visualization analogous to C for a spring relaxation reconstruction. (F) Radial profile analogous to D for the spring relaxation reconstruction approach.

## 5 Approach Variations

Due to the high number of non-information bearing pairing events that is most likely to occur with the above technique, i.e.barcodes of one polony pairing with neighboring barcodes of the same polony, we propose some advanced variations on the basic principle. One approach would be to use a bipartite network formation. The bridge amplification approach to polony generation, for example, leaves the possibility of having two species of independent primers on the surface and the formation of essentially two interpenetrating/overlapping and indepedentently saturated polony surfaces. By introducing crosslinking that bridges the two networks, a bipartite network would be formed where essentially every pairing event would be information bearing, since each cross link would necessarily have two distinct barcodes (one of each polony species). Another possible approach would be series growth of polonies. In the basic concept presented in previous sections, a primer of uniform sequence is assumed, however generation of a saturated layer of polonies that could then be used as primers for a subsequent polony generation step would then result in an overlapping of every 2nd-layer polony with multiple 1st-layer polony.

## 6 Conclusion

PARSIFT is a concept for microscopic image construction based on the encoding of spatial information into the format of DNA bases that can be reconstructed after sequencing. Here we have shown a mathematical basis for the transmission of spatial information via topology that stems from the relationship of Euclidean neighborhood with that of topological geodesic neighborhood, the former of which can be inferred from the latter. We have demonstrated an in silico proof of concept by constructing a pipeline for taking decoupled edge data, generated from simulated polony distributions, that are then reassembled into a topological network and embedded in a Euclidean plane, resuming much of the spatial characteristics of the original seed distribution. We saw that global distortion is largely absent, and that local distortions decay with increasing polony density. We hold that this framework and pipeline for reconstruction could be exploited for image analysis of micro- and nano-scale surfaces with molecular libraries of potentially very high multiplicity and with throughput automated in a way that would not be possible with most optical approaches.

